# Differential Astrocyte-supplied NMDAR Co-Agonist for CA1 versus Dentate Gyrus Long-term Potentiation

**DOI:** 10.1101/2025.05.05.652314

**Authors:** Shruthi Sateesh, Wickliffe C Abraham

## Abstract

In the hippocampus, there is a region and synapse-specific N-methyl-D-aspartate receptor (NMDAR) co-agonist preference for induction of long-term potentiation (LTP). Schaffer collateral (SC)-CA1 synapses, enriched in GluN2A-containing NMDARs, favor D-serine, while medial perforant path (MPP) to dentate gyrus (DG) synapses that are rich in GluN2B-containing NMDARs prefer glycine for LTP induction. This study investigated the role of astrocytes in providing these co-agonists. We confirmed in rat hippocampal slices that exogenous D-serine (10 µM) is sufficient to restore LTP at SC-CA1 synapses blocked under astrocyte calcium (Ca^2+^) -clamp conditions, consistent with previous findings. However, exogenous glycine (10 μM) also rescued the LTP. In contrast, at MPP-DG synapses, 100 µM exogenous glycine, but not 10 µM nor 100 µM D-serine, restored the LTP blocked by astrocyte Ca^2+^-clamping. Our findings support the view that, as for serine in CA1, astrocytes are the cellular source of the glycine required for LTP induction at MPP-DG synapses.

---

Long-term potentiation (LTP), a persistent strengthening of synaptic transmission, is a widely accepted cellular correlate of learning and memory (Abraham et al., 2019; Bliss & Collingridge, 1993). This process is fundamentally dependent on the activation of N-methyl-D-aspartate receptors (NMDARs). NMDAR activation requires the simultaneous binding of glutamate to GluN2 subunits and a co-agonist binding to the glycine site on GluN1 subunits, with glycine and D-serine being the two prominent examples. Glycine plays a role in enabling glutamate-mediated NMDAR activation and subsequent LTP induction broadly across the hippocampus (Abe et al., 1990; Akio et al., 1991; Johnson & Ascher, 1987).

In contrast to glycine, D-serine, has been shown to be particularly significant for LTP induction in hippocampal CA1 (Mothet et al., 2000), where LTP induction at the Schaffer collateral (SC) synapses relies heavily on the presence of D-serine (Han et al., 2015; Mothet et al., 2006). Notably, astrocytes were identified as the source of D-serine, as clamping of internal calcium (Ca^2+^) in individual CA1 astrocytes blocked LTP induction at nearby excitatory synapses by decreasing the occupancy of the NMDAR co-agonist sites. This LTP blockade was reversed by exogenous bath application of D-serine, either through its direct release (Henneberger et al., 2010) or indirectly through the release of L-serine, which is subsequently converted to D-serine by neurons via serine racemase (Neame et al., 2019). Interestingly, this astrocytic source of D-serine metaplastically regulates LTP in the supraoptic nucleus (Panatier et al., 2006). On the other hand, glycine can also play a role in LTP induction in the CA1 region of the hippocampus (Shahi et al., 1993; Zhang et al., 2014), with co-agonist preference depending in part on the developmental age of the hippocampus (Le Bail et al., 2015).

The identity of the NMDAR co-agonist in the hippocampus correlates with the GluN2 subunit composition of postsynaptic NMDARs and the maturation of the tripartite synapse (Le Bail et al., 2015; Papouin et al., 2012). Previous studies utilizing enzymatic scavengers and pharmacological inhibition of D-amino acid oxidase have revealed a distinct compartmentalization: D-serine is the preferred co-agonist at mature SC-CA1 synapses, whereas glycine predominantly drives LTP at medial perforant path (MPP)-dentate gyrus (DG) synapses (Le Bail et al., 2015; Musleh et al., 1997). However, whether astrocytes are the main source of glycine governing LTP at MPP-DG synapses, as they are for D-serine in CA1 (Henneberger et al., 2010), remains unknown. This study addressed this gap by investigating whether Ca^2+^-dependent release of glycine from astrocytes plays a preferential role in LTP induction at MPP synapses in the middle molecular layer (MML) of the DG.

Young adult (6–8-week-old) male Sprague-Dawley rats were obtained from colonies maintained at the University of Otago Breeding Station. All animals were maintained at an ambient temperature of 24 °C ± 1°C and under a standard 12-hour light/dark cycle. Animals were group-housed, with no more than 4 per cage, and had free access to food and water. All experiments were conducted in accordance with New Zealand animal welfare legislation, and ethical approval was gained from the University of Otago Animal Ethics Committee (AUP #18-51 and #19-42). For electrophysiological recordings, animals were deeply anesthetized with ketamine (100 mg/kg, i.p.), and then the hippocampus was dissected, CA3 removed, and 400 μm slices prepared in ice-cold, carbogenated sucrose dissection solution (in mM: 210 sucrose, 26 NaHCO_3_, 2.5 KCl, 1.25 NaH_2_PO_4_, 0.5 CaCl_2_, 3 MgCl_2_, 20 D-glucose). Slices were incubated in carbogenated artificial cerebrospinal fluid (ACSF; in mM: 124 NaCl, 3.2 KCl, 1.25 NaH_2_PO_4_, 26 NaHCO_3_, 2.5 CaCl_2_, 1.3 MgCl_2_, 10 D-glucose) at 32°C with 0.5 μM sulforhodamine 101 (SR101-Sigma, S7635, **Fig. 1A**) for astrocyte labelling (Nimmerjahn & Helmchen, 2012), then transferred to room temperature ACSF. After at least 2 hours, slices were transferred to the recording chamber and perfused with ACSF at 32.5°C and a flow rate of 2.5-3 ml/min. DIC microscopy with a 40x objective (Olympus BX51WI) was used for imaging. SR101-positive astrocytes in the stratum radiatum (SR) of CA1 and MML of DG using epifluorescence. Whole-cell patch-clamp recordings were performed on individual astrocytes (**Fig. 1B**). Micropipettes (2.5-4.5 MΩ) were filled with a potassium-based intracellular solution (mM: KMeSO_4_ 130, HEPES 10, Na_2_-ATP 4, Na_2_-GTP 0.4, Na_2_-phosphocreatine 4, MgCl_2_, pH 7.2, 295–300 mOsM). For Ca^2+^ clamping, the intracellular solution contained 0.45 mM EGTA and 0.14 mM CaCl_2_ to maintain Ca^2+^ at 50–80 nM (Henneberger et al., 2010). Under clamped conditions, the extracellular ACSF included either D-serine (Sigma, S4250) or glycine (Sigma, G7126). Astrocytes were patched in either CA1 SR or the DG MML and identified visually by their resting membrane potential (-80 mV) and by their linear current-voltage relationships (**Fig. 1C**). Local field potential recordings were made through the astrocyte membrane in current-clamp mode. Synaptic potentials were evoked by stimulating the fibers in SR or MML using programmable stimulators. The recording electrode was placed ∼350 μm from the stimulating electrode, and recordings were digitized at 10 kHz. Synaptic potentials recorded from the patched astrocytes (AfEPSPs) were smaller than typical extracellularly recorded fEPSPs (**Fig. 1D**, Henneberger et al., 2010). For LTP induction in SR, two trains of theta-burst stimulation (TBS) were applied. Each train consisted of 10 bursts of 5 pulses at 100 Hz, with a pulse duration of 100 µs and 200 ms between bursts, at a fixed current strength of 75 µA. For LTP induction in MPP synapses in MML, four trains of TBS were applied, with 30 s between each train. Each train consisted of 10 bursts and 10 pulses at 100 Hz, with a pulse duration of 200 µs and 200 ms between bursts, at a fixed current strength of 55 µA, in the presence of 0.2 µM gabazine (SR 95532 hydrobromide, HB0901). To qualitatively confirm SR stimulation in CA1 or MPP stimulation in the DG, paired-pulse tests were performed by delivering two pulses 50 ms apart (**Fig. 1D,E**). Data were acquired using Clampex version 10.7 and analysed using Clampfit version 10.7 (Molecular Devices). The initial slopes of the fEPSPs were measured off-line and expressed as a percentage of the baseline value obtained by averaging the fEPSP slope for the 10 min preceding TBS. All data are expressed as mean ± SEM. Statistical comparisons were performed using GraphPad Prism software 10.4.2, using one-way ANOVA. For multiple comparisons, analysis was performed using Uncorrected Fisher’s LSD post-hoc tests. The data for each group were pooled across animals, with individual slices being the experimental unit of analysis (n’s), and group sizes varied between n = 3-6.

**Figure 1:**
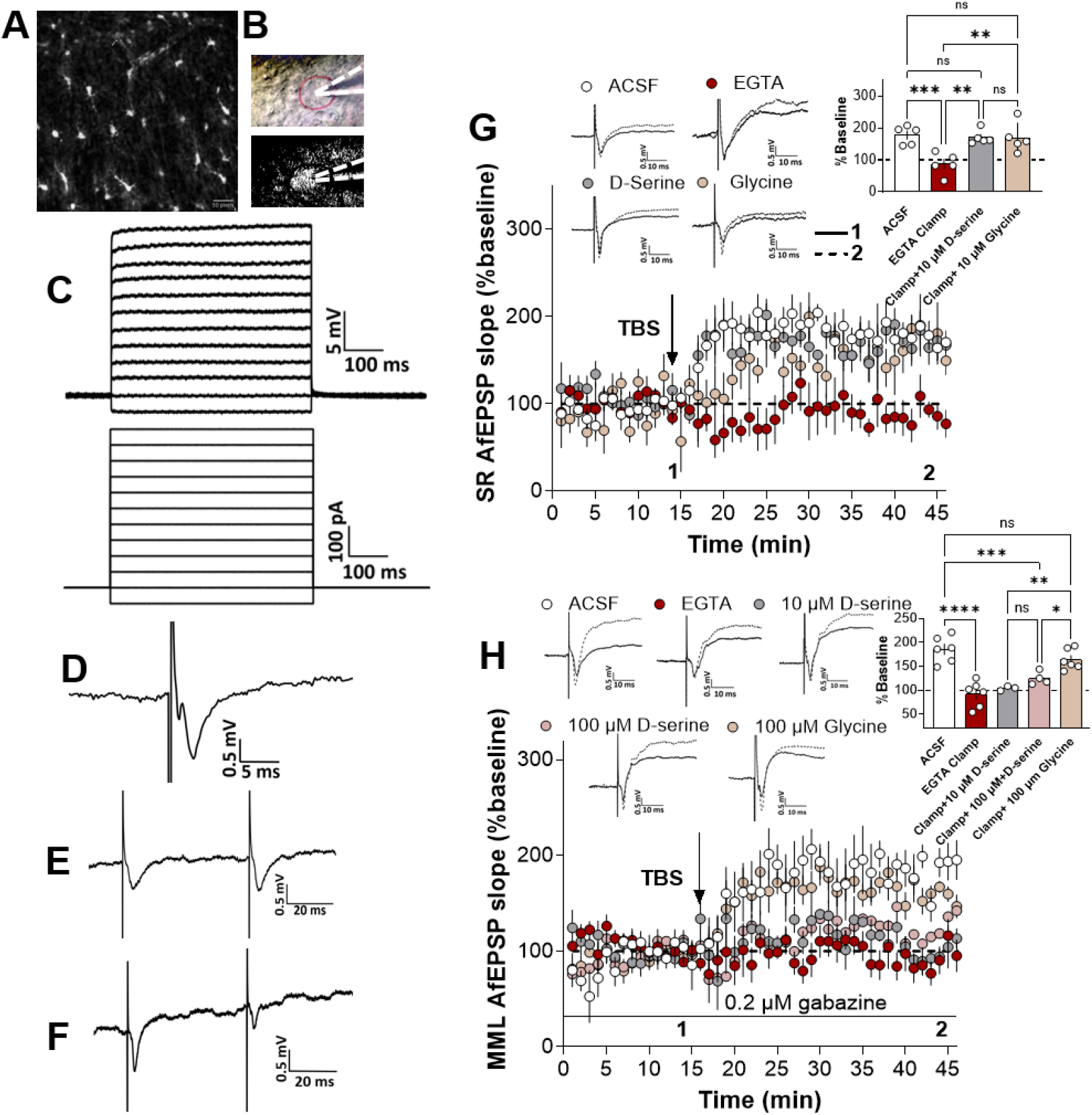
(**A)** Representative confocal image of astrocytes labelled with SR101 using a 10x water immersion objective. (**B**) Representative images showing differential interference contrast (DIC) and corresponding fluorescence image of an SR 101 patched astrocyte in MML (glass pipette shown in white dotted line) using a 40x water immersion objective lens. (**C**) Schematic of 100 pA current steps used to generate V_m_ deflections and the corresponding traces of current-voltage relation from a single whole-cell patch-clamped astrocyte. (**D**) Representative field response in MML recorded through an astrocytic membrane (AfEPSP) following test pulse stimulation of the MPP inputs.(**E,F**) Representative average waveform of 4 sweeps showing either paired-pulse facilitation or paired-pulse depression as recorded through the astrocyte membrane when two pulses were delivered at SC-CA1(**E**) and MPP-DG (**F**) synapses at a 50-ms inter-pulse interval. (**G**) Clamping astrocytic Ca^2+^ with EGTA blocked LTP at nearby SC synapses in SR of CA1 in a D-serine (10 μM) as well as glycine (10 μM)-dependent manner. (**H**) Clamping astrocytic Ca^2+^ blocks LTP at nearby MMP synapses in MML of DG in a specifically glycine-dependent manner. LTP was measured in the presence and absence of clamping and by supplying either 10 μM, 100 μM D-serine, or 100 μM glycine under Ca^2+^ clamped conditions. (**G,H**) *Inset:* Bar graphs summarising the LTP measured via the whole-cell patch-clamped astrocyte in the presence and absence of Ca^2+^ clamping. Representative waveforms are an average of 10 synaptic responses prior to LTP induction (1) and at the conclusion of the experiment (2). The arrow indicates the timing of LTP induction. All data presented as mean ± SEM; ns, p > 0.05; *, p < 0.05; **, p < 0.01; ***, p < 0.001; ****, p < 0.0001.

In the CA1 SR, two trains of TBS induced robust LTP in control conditions (no Ca^2+^-clamping: 175.4 ± 14.5%, n=5). As previously reported by Henneberger et al (2010), clamping astrocytic Ca^2+^ with EGTA completely blocked this LTP (88.1 ± 14.3%, n=5; p= 0.0007 compared to controls, **Fig. 1G**), but bath application of 10 µM D-serine throughout the experiment rescued the LTP (170.3 ± 8.8%, n=5; p= 0.74, **Fig. 1G**). Interestingly, bath application of 10 µM glycine also restored the LTP (Glycine: 169.4 ± 20.7%, n=5; p= 0.70, **Fig. 1G**). On the other hand, the equality of D-serine and glycine in rescuing LTP was not observed in the MML of DG. Here, LTP was again blocked by the astrocyte Ca^2+^-clamp (91.8± 11.1%, n=6; p<0.0001, **Fig. 1H**) compared to control non-clamp conditions (185.3 ± 11.1%, n=6, **Fig. 1H**), but neither 10 µM D-serine nor the 10-fold higher concentration of 100 µM restored LTP (10 µM: 103.8 ± 2.8%, n=3, p <0.0001; 100 µM : 124.4 ± 6.7%, n=4, p= 0.0005, **Fig. 1H**). In contrast, 100 µM glycine during Ca^2+^-clamping readily restored LTP to near control levels (163.7 ± 8.8%; p= 0.11, **Fig. 1H**). These findings indicate that NMDAR-dependent LTP at the MPP synapses in the DG is particularly dependent on Ca^2+^-dependent glycine release from astrocytes.

Astrocytes can produce both D-serine and glycine function as co-agonists at NMDARs that are crucial for LTP. The synthesis of D-serine primarily occurs from L-serine, a process involving serine racemase, with L-serine itself derived from phosphoglycerate dehydrogenase metabolism of glucose within astrocytes (Mothet et al., 2015; Neame et al., 2019). Glycine is also synthesized within astrocytes (Singer & Yee, 2024; Tripodi et al., 2023; Verleysdonk et al., 1999) and the release of both co-agonists is activity-dependent, enabling NMDAR function, particularly during periods of heightened activity (Abreu et al., 2023; Li et al., 2009; Li et al., 2013). This study highlights the critical role of astrocytes in regulating LTP in the hippocampus through differential involvement of the NMDAR co-agonists D-serine and glycine in distinct hippocampal subregions. Specifically, our findings demonstrate that Ca^2+^-dependent mechanisms within astrocytes are essential for the release of glycine that is critical for LTP induction at MPP-DG synapses, mirroring previous findings regarding D-serine release in the CA1 region (Henneberger et al., 2010).

Previous work indicates a regional specialization of co-agonist function: D-serine is particularly important for LTP at SC synapses in CA1, while glycine predominates at MPP-DG synapses. This aligns well with developmental changes in NMDAR subunit composition, with CA1 synapses transitioning to GluN2A-containing NMDARs, which have a higher affinity for D-serine (Le Bail et al., 2015), whereas MPP-DG synapses retain more GluN2B-containing NMDARs, which favor glycine (Ferreira et al., 2017; Le Bail et al., 2015). Supporting the importance of endogenous glycine at MPP synapses, studies have shown that blocking glycine with antagonists such as 7-chlorokynurenate impairs LTP in the DG (Akio et al., 1991).

While the co-agonist preference differences between CA1 and DG are established, the new findings emphasize the astrocytes as the source of these co-agonists. Specifically, we replicated Henneberger et al. (2010) by first confirming that D-serine derived from astrocytes is important for LTP at SC-CA1 synapses. In addition, we showed that glycine at the same concentration is also capable of rescuing CA1 LTP under Ca^2+^-clamp conditions. Note, however, that at elevated concentrations in CA1, glycine can contribute to long-term depression (LTD) through the activation of glycine receptors (Chen et al., 2011; Zhang et al., 2014), and at very high concentrations, it can induce excitotoxicity via excessive NMDAR activation (McNamara & Dingledine, 1990; Newell et al., 1997).

D-serine and glycine not only act as co-agonists but also influence NMDAR trafficking, thereby providing a different mechanism for modulating synaptic plasticity. D-serine in particular has been shown to affect the synaptic content of GluN2B-containing NMDARs (Ferreira et al., 2017; Papouin et al., 2012). This suggests that the relative availability of these co-agonists can dynamically regulate NMDAR subunit composition at the synapse, impacting synaptic strength. Endogenous concentrations of D-serine and glycine in the hippocampus also play a crucial role in synaptic plasticity. While both amino acids increase from the neonatal period to adulthood, the increase in D-serine is more pronounced in CA1, partly explaining its role in restoring LTP under Ca^2+^-clamped conditions (Le Bail et al., 2015). Notably, the same study showed a higher concentration of endogenous glycine compared to D-serine in adult DG (Le Bail et al., 2015). These regional differences in endogenous co-agonist concentrations no doubt contribute to the distinct roles of D-serine and glycine in modulating synaptic plasticity across hippocampal sub-regions.

In conclusion, this study provides evidence that astrocytes are a critical source of glycine for LTP induction at MPP-DG synapses, highlighting the differential NMDAR co-agonist requirements for LTP in the hippocampus and the essential but differential roles of astrocyte-neuron interactions in supporting hippocampal synaptic plasticity.

## Acknowledgements

This research was funded by grants from the Health Research Council of New Zealand (#18/245) to W.C.A, and (#22/177) to W.C.A and S.S. Additionally, S.S. received a New Zealand International Doctoral Scholarship, a Roche Hanns Möhler Doctoral Scholarship, and a Brain Research New Zealand research grant.

## Notes

### Competing Interest Statement

The authors have declared no competing interest.

